# Accounting for Cellular-Level Variation in Lysis: Implications for Virus-Host Dynamics

**DOI:** 10.1101/2024.05.04.592515

**Authors:** Marian Dominguez-Mirazo, Jeremy D. Harris, David Demory, Joshua S. Weitz

**Affiliations:** School of Biological Sciences, Georgia Institute of Technology, Atlanta, GA, USA; Interdisciplinary Graduate Program in Quantitative Biosciences, Georgia Institute of Technology, Atlanta, Georgia, USA; Department of Mathematics, Rose-Hulman Institute of Technology, Terre Haute, Indiana, USA; CNRS, Sorbonne Université, USR3579 Laboratoire de Biodiversité et Biotechnologies Microbiennes (LBBM), Observatoire Océanologique, Banyuls-sur-Mer, F-66650, France; Department of Biology, University of Maryland, College Park, Maryland, USA; Department of Physics, University of Maryland, College Park, Maryland, USA; Institut de Biologie, École Normale Supérieure, Paris, France

## Abstract

Viral impacts on microbial populations depend on interaction phenotypes - including viral traits spanning adsorption rate, latent period, and burst size. The latent period is a key viral trait in lytic infections. Defined as the time from viral adsorption to viral progeny release, the latent period of bacteriophage is conventionally inferred via one-step growth curves in which the accumulation of free virus is measured over time in a population of infected cells. Developed more than 80 years ago, one-step growth curves do not account for cellular-level variability in the timing of lysis, potentially biasing inference of viral traits. Here, we use nonlinear dynamical models to understand how individual-level variation of the latent period impacts virus-host dynamics. Our modeling approach shows that inference of latent period via one-step growth curves is systematically biased - generating estimates of shorter latent periods than the underlying population-level mean. The bias arises because variability in lysis timing at the cellular level leads to a fraction of early burst events which are interpreted, artefactually, as an earlier mean time of viral release. We develop a computational framework to estimate latent period variability from joint measurements of host and free virus populations. Our computational framework recovers both the mean and variance of the latent period within simulated infections including realistic measurement noise. This work suggests that reframing the latent period as a distribution to account for variability in the population will improve the study of viral traits and their role in shaping microbial populations.

**Importance:** Quantifying viral traits – including the adsorption rate, burst size, and latent period – is critical to characterize viral infection dynamics and to develop predictive models of viral impacts across scales from cells to ecosystems. Here, we revisit the gold standard of viral trait estimation – the one-step growth curve – to assess the extent to which assumptions at the core of viral infection dynamics lead to ongoing and systematic biases in inferences of viral traits. We show that latent period estimates obtained via one-step growth curves systematically under-estimate the mean latent period and, in turn, over-estimate the rate of viral killing at population scales. By explicitly incorporating trait variability into a dynamical inference framework that leverages both virus and host time series we provide a practical route to improve estimates of the mean and variance of viral traits across diverse virus-microbe systems.

## 1. Introduction

Viruses have a profound impact on microbial populations through the modulation of population dynamics, eco-evolutionary dynamics, community structure, and ecosystem function [1–6]. Our understanding of the ecological interactions between viruses and microbes depends on the study of viral life history traits that characterize the viral life cycle, such as adsorption rate, latent period, and burst size [7, 8]. During lytic infections, viral adsorption to the host cell is followed by synthesis where the host machinery produces new viral particles. Once assembled, viral particles burst from the cell and can infect new target cells. The time to complete a viral cycle, from virus adsorption to cell burst, is termed the latent period [7]. The latent period varies across taxa [8] and can be influenced by environmental factors such as resource availability and host physiological state [9–12]. This key trait affects the population dynamics of virus-microbe pairs, with strong consequences for virus fitness and host population fate [13, 14].

The one-step growth curve, introduced by Ellis and Delbrück in 1939 [15], is an experimental setup used to infer key viral traits from monitoring viral infection of a microbial population within a single round of infection. Typically, the one-step growth curve has the following experimental protocol: First, cells are inoculated with virus under conditions that allow for adsorption to take place without host reproduction or viral replication. Then, unadsorbed viruses are removed to allow for a single round of infection. Finally, free viruses are measured at several time points. From these observations, the latent period is typically reported as the time of first visible burst (Figure 1), *i.e*., when free virus concentration first increases [15, 16]. Alternatively, the midpoint of the rise might be used to report the latent period [8, 12]. Despite widespread use, the first visible burst can lead to biases in estimates of the average latent period in the presence of cellular-level variability in lysis, as it captures the combined influence of early adsorption with early cell lysis events [15, 17–20].

**Figure 1:**
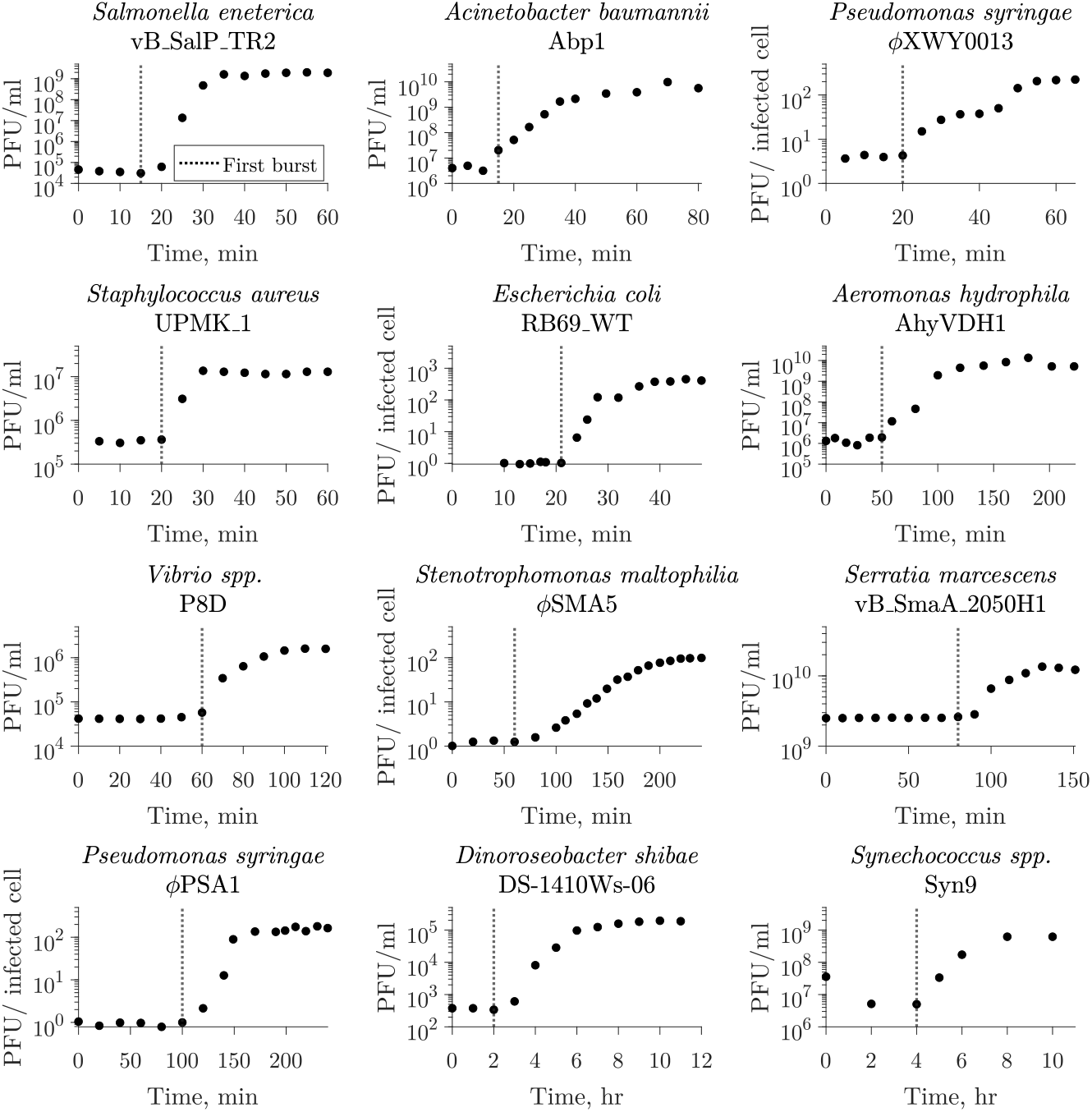
The one-step growth curve protocol for inferring lysis timing. The one-step growth curve is used to estimate burst size and latent period by observing a single round of infection. In such experiments, virus is added to a microbial population and left to adsorb until the majority of the cells are infected. The population is diluted or viruses are removed to prevent the occurrence of new infections. From this point, plaque-forming units (PFUs) are measured over time. The time of first visible burst, when PFU counts starts to increase due to cell lysis and viral progeny release, is commonly reported as the latent period [18, 21]. The latent period of multiple virus-microbe pairs has been characterized using this method. Here, we show examples of different microbe-virus pairs where the time of first burst was reported as the latent period ranging from 15 min to 4 hrs. The dotted line represents the reported value in the corresponding study. The list of data sources [10, 22–32] is available in Table S1.

Indeed, it is already well understood that even when an infected cell population is perfectly synchronized, *i.e*., ensuring adsorption synchronization, individual cells may burst at different times. This variation could be due to differences in host phenotype at the time of infection, *e.g*., physiological states [12, 17, 19, 33, 34], or due to stochasticity at the cellular level [20, 35–37]. For instance, variability in the latent period of genetically identical populations has been found in multiple strains of *λ* phage through single cell analysis [14, 35]. The majority of this variation was accounted for by processes that regulate the production of holins that trigger lysis by forming holes in the inner membrane and allowing endolysin into the periplasmic space. Variation in holin genotypes can lead to different mean latent periods suggesting a (partially) viral-encoded origin of cell-to-cell variability in latent period [35, 36]. In addition, recent work has characterized heterogeneity in timing for a lytic phage for the first time, showing that variability in lysis time may provide fitness advantages to phage populations [20].

Here, we explore the effect of latent period variability on microbe-virus dynamics and its impact on current approaches to infer the latent period in practice. Our findings reveal that the presence of latent period variability results in systematic biases that can lead to the underestimation of latent periods in the one-step growth curve used for viral trait estimation. Hence, established methods infer a more rapid lysis process from population measurements than what is likely to occur when accounting for cellular-level heterogeneity. Instead, by integrating cellular-level heterogeneity in lysis timing in an explicit virus-microbe infection model we are able to estimate both the mean and variability of lysis timing from population-level data. As we show, expanding current protocols to include measurements of both virus and host abundances via the use of ‘multi-cycle response curves’ provides a route to improved estimates of viral traits and their variability, essential to quantify the impact of viruses on microbial populations in both ecological and therapeutic contexts.

## 2. Results

### 2.1 Latent period variability impacts one-step growth curves

We use a coupled system of nonlinear differential equations to model interactions between a population of microbial cells and a lytically infecting viral population (Methods, Figure S1). The model considers a microbial population where infected cells burst at different times following an Erlang distribution described by the mean latent period and the coefficient of variation (CV). We perform simulations of one-step growth curves to elucidate the impact of individual-level variation on conventional methods of viral trait estimation (Figure 2). As described in the methods, we replicate a classic one-step growth curve protocol where phage is added to a bacterial population in exponential growth at a MOI of 0.01, the system is diluted to reduce new infections, and the accumulation of free viruses is measured over time (Figure 2A).

**Figure 2:**
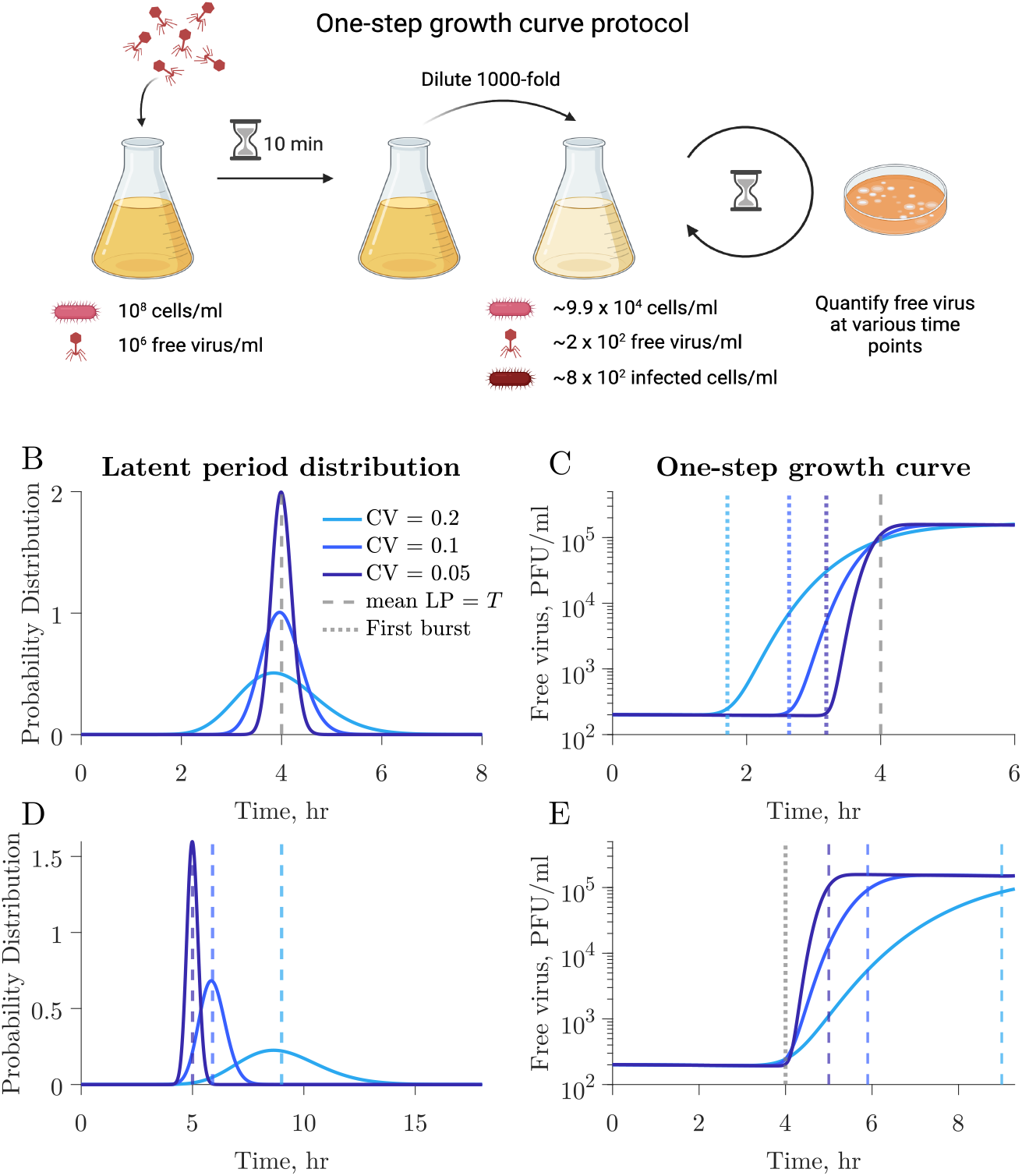
The latent period distribution connects individual variation to population-level microbe-virus dynamics. **A)** Using our microbe-virus dynamical model, we simulated one-step growth curves following standard protocols, as described in Methods. **B)** Populations in each simulation have the same traits (Table 1), *i.e*., microbial growth rate, carrying capacity, adsorption rate, burst size, and latent period (LP) mean (4 hr, dashed gray line), and only differ in the CV of the latent period distribution which varies across simulations with larger CV depicted in lighter shades of blue. **C)** The one-step growth curves for the different simulations show different free virus dynamics. The standard estimates of the latent period, as measured by the time of first burst (dotted lines) vary across simulations. Latent period variability affects the one-step growth curves for otherwise identical populations. **D)** In this set of simulations, the populations have the same traits (Table 1), *i.e*., microbial growth rate, carrying capacity, adsorption rate, burst size, but differ in latent period distributions with varying latent period mean and CV. **E)** Systems with visibly different latent period distributions can result in similar first burst estimates derived from one-step growth curves.

Figure 2B,C shows three different simulations in which all host and viral traits are consistent across scenarios including the average latent period, except for the latent period CV, which varies across 0.05, 0.1, to 0.2 (see Table 1 for complete specification of parameter values). The resulting one-step growth curves are visibly different between simulations. Furthermore, the time of first visible burst, which is conventionally used as a proxy for the average latent period (Figure 1), varies greatly for the different simulations despite them having the same mean latent period. This also applies to other features of the curve, *e.g*., the midpoint of the rise. Specifically, in wider distributions (larger CV) the first visible burst occurs earlier because of a higher proportion of cells lysing at shorter times. Hence, the time of first burst leads to a biased estimate of shorter latent periods than the true, underlying latent period for all simulated distributions, including distributions with small CV. These results indicate that reporting the time of first burst as latent period is a misleading conflation: doing so systematically underestimates the population mean and this bias worsens with increasing cellular-level variability.

**Table 1:** Model parameter values for one-step growth curves. We simulate one-step growth curves in Figure 2 using the parameter set and initial conditions specified here. The CV and lysis rate (*η*) for each simulation are specified in the corresponding figure. The lysis rate is represented as the mean latent period (*T* = 1*/η*).

| Parameter | Value | Unit |
| --- | --- | --- |
| $\mu$, growth rate | 0.1 | hr<sup>-1</sup> |
| $K$, carrying capacity | $1 \times 10^9$ | CFU/ml |
| $\phi$, adsorption rate | $1 \times 10^{-7}$ | ml/(CFU×hr) |
| $\beta$, burst size | 200 | |
| $\eta$, lysis rate | * | hr<sup>-1</sup> |
| CV, coefficient of variation | * |  |
| Initial condition | Value | Unit |
| $S_0$, initial microbial density | $10^8$ | CFU/ml |
| $V_0$, initial free virus density | $10^6$ | PFU/ml |

\*Value specified in the corresponding section

We further illustrate the influence of latent period variability on key features of the one-step growth curve by comparing three simulations with varying latent period distributions, each characterized by distinct mean and CV (Figure 2D). Host and viral traits, excluding latent period-associated traits, remain the same across all simulations (see Table 1 for parameter values). Despite distinct underlying latent period distributions, including differences in the mean latent period, the simulated one-step growth curves consistently result in the same time of first burst (Figure 2E, dotted line).

Based on these observations we revisited data from [13, 35] to compare the differences between population and cellular-level measurements of lysis timing. In these studies the lysis time, *i.e*., the time to lysis after prophage induction, was measured using population-scale turbidity assays [13] and single-cell lysis event microscopy observations [35] of isogenic *λ* lysogens with mutations in holin and antiholin coding genes. We observe that the lysis time is systematically shorter in population-scale turbidity assays when compared to the mean lysis time obtained from quantification of single-cell lysis events (Figure 3). The discrepancy between measurements follows a trend similar to that predicted by theory, especially in mutants where the antiholin gene expression is abolished (Figure S2). Although variations in bacterial treatment could contribute to the observed differences, the disparity between measurements at the population scale and those at the single-cell level may indicate that underestimation of life-history traits regarding timing in lysis may be a generic feature of population-level protocols, and relevant to both temperate and lytic-only viruses.

**Figure 3:**
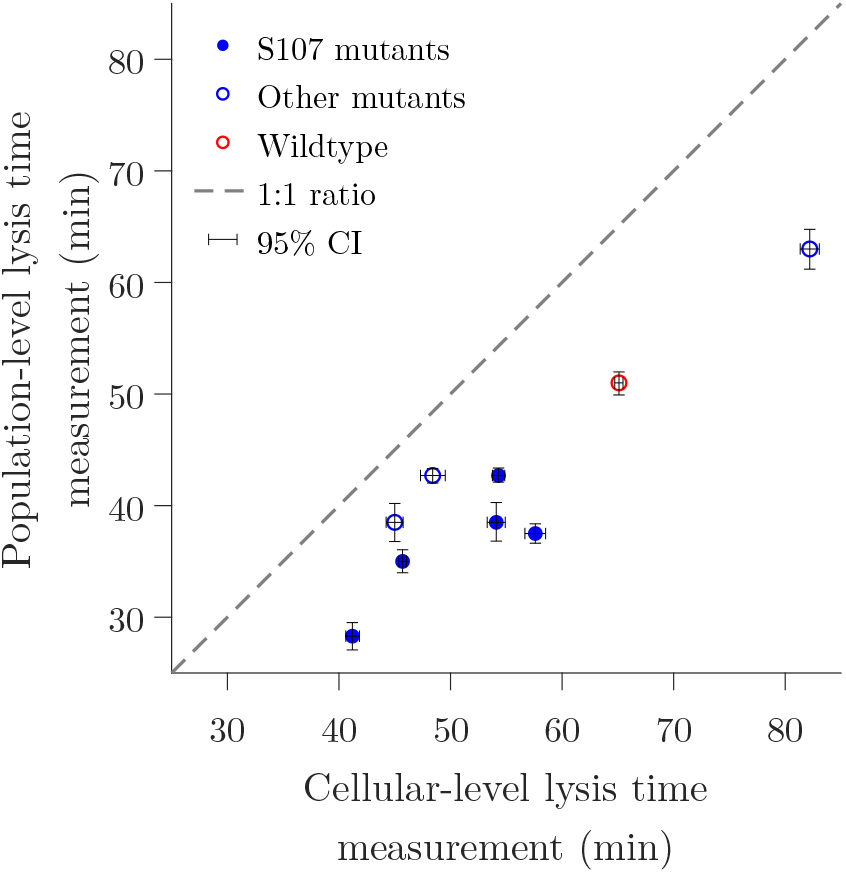
Population-level measurements systematically underestimate the lysis time in *λ* phage lysogens. The measured lysis time, *i.e*., time to lysis after prophage induction, of *λ* lysogens is systematically shorter in population-scale turbidity assays [13] when compared to the mean lysis time obtained from microscope observations of single-cell lysis events [35]. The lysogens are isogenic with mutations in holin and antiholin coding genes that result in changes in lysis time. Solid circles represent S107 mutants where antiholin expression is abolished. The dashed line shows a 1 to 1 relationship. Confidence intervals were calculated by multiple resampling using experimental mean and standard deviation assuming normality, as explained in [38]. Data recovered from [13, 35] is available in Table S2.

### 2.2 Free-virus and host temporal dynamics provide better resolution than one-step growth curves to discern between latent period distributions

Next, we set out to explore the link between viral population dynamics arising from one-step growth curve protocols and the shape of the underlying latent period distribution. We systematically varied both the mean and CV of the latent period distribution. In each case, we simulated the one-step growth curve protocol and compared the resulting viral population dynamics to a reference (*i.e*., the true, but unknown mean and CV of the viral-host pair). For example, given a reference latent period of 10 hrs and CV 0.25 we calculated the sum of relative errors in viral population dynamics for each combination of latent period and CV (Figure 4A). Growth curves resulting from combinations featuring a higher mean and larger CV will resemble those with a smaller mean and smaller CV (Figure 4A). This, coupled with experimental uncertainty, results in a space of latent period distributions difficult to distinguish from each other when comparing one-step growth curves (Figure 4B,C). These results suggest that one-step growth curve protocols lack the resolution needed for accurate characterization of latent period distributions.

**Figure 4:**
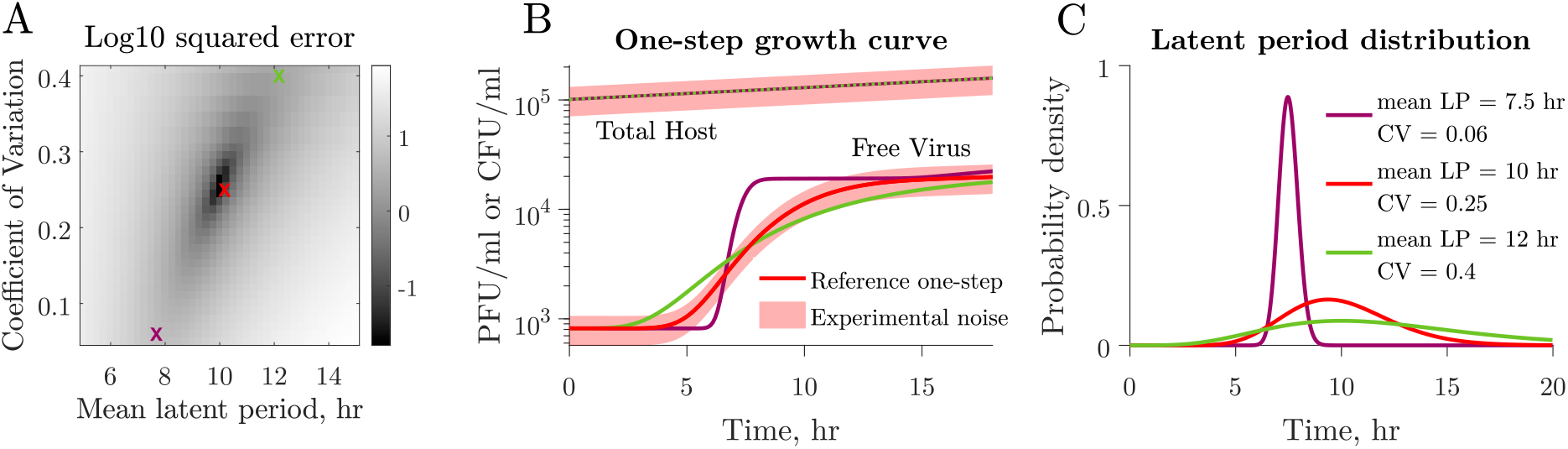
Latent period distribution identifiability when using one-step growth curves. **A)** Simulated one-step growth curves (see Methods) obtained from microbe-virus pairs with different underlying latent period distributions can resemble each other. When we compare a reference curve (red cross) to curves obtained from systems with different distributions, we observe that curves that resemble the reference the most are found along an ascending slope. These correspond to combinations of larger mean, larger CV (green cross) or smaller mean, smaller CV (purple cross). **B)** Example of different combinations of latent period mean and CV that produce similar curves. One-step growth curves become harder to differentiate when taking experimental noise into account. Changes in host density, represented by colony-forming units per volume unit, resulting from viral lysis in a single cycle of infection are insignificant owing to the low MOI utilized in protocols. **C)** Corresponding latent period distributions for panels A and B. All non-latent period traits, *i.e*., microbial growth rate, carrying capacity, adsorption rate, burst size, are the same across simulations (Table 2).

As an alternative, we propose the use of a ‘multi-cycle response curve’ protocol. In this protocol, phage are mixed with a susceptible bacterial population at low MOI. Next, both free virus and host cells are quantified at various time points after phage addition (Figure 5). In contrast to the one-step growth curve simulations, there is neither an incubation period nor a subsequent dilution. While some combinations of mean latent period and CV result in similar host and viral population dynamics (Figure 4), the corresponding host population dynamics for these combinations are markedly different from each other (Figure 5C). These observations imply that population-level temporal dynamics data of hosts and viruses have the potential to be used to predict individual-level variation in viral traits.

**Table 2:**
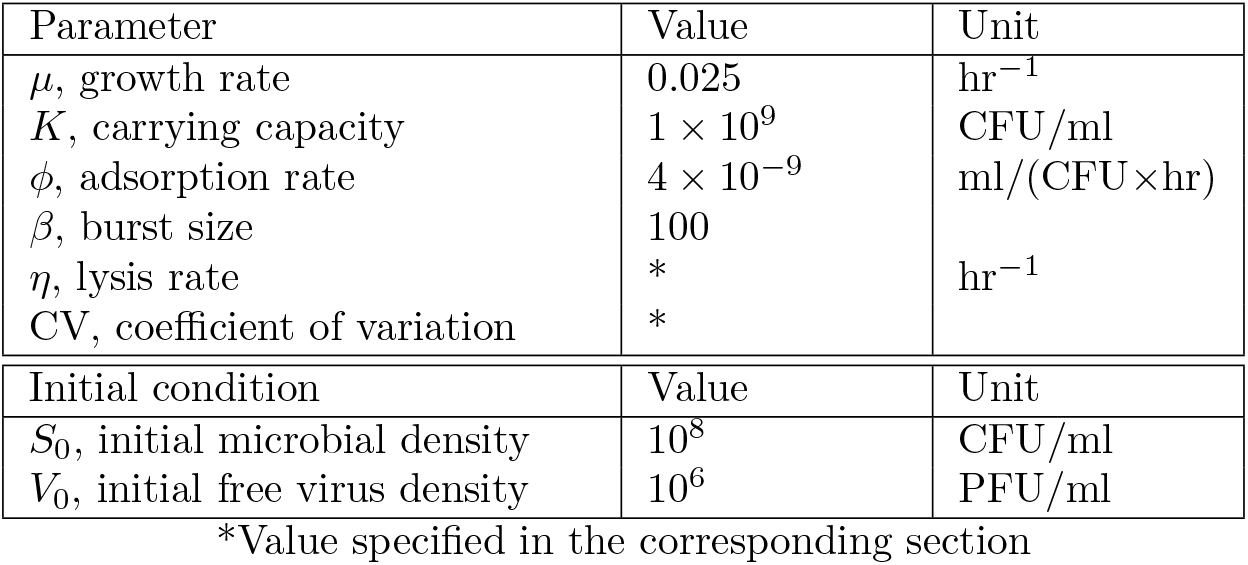
Model parameter values for one-step growth curves and multi-cycle response curves. We simulate one-step growth curves and the corresponding multi-cycle response curves in Figure 4 and Figure 5 using the parameter set and initial conditions specified here, and obtained from [41]. The CV and lysis rate (*η*) for each simulation are specified in the corresponding figure. The lysis rate is represented as the mean latent period (*T* = 1*/η*).

| Parameter | Value | Unit |
| --- | --- | --- |
| $\mu$, growth rate | 0.025 | hr<sup>-1</sup> |
| $K$, carrying capacity | $1 \times 10^9$ | CFU/ml |
| $\phi$, adsorption rate | $4 \times 10^{-9}$ | ml/(CFU×hr) |
| $\beta$, burst size | 100 | |
| $\eta$, lysis rate | * | hr<sup>-1</sup> |
| CV, coefficient of variation | * |  |
| Initial condition | Value | Unit |
| $S_0$, initial microbial density | $10^8$ | CFU/ml |
| $V_0$, initial free virus density | $10^6$ | PFU/ml |

\*Value specified in the corresponding section

**Figure 5:**
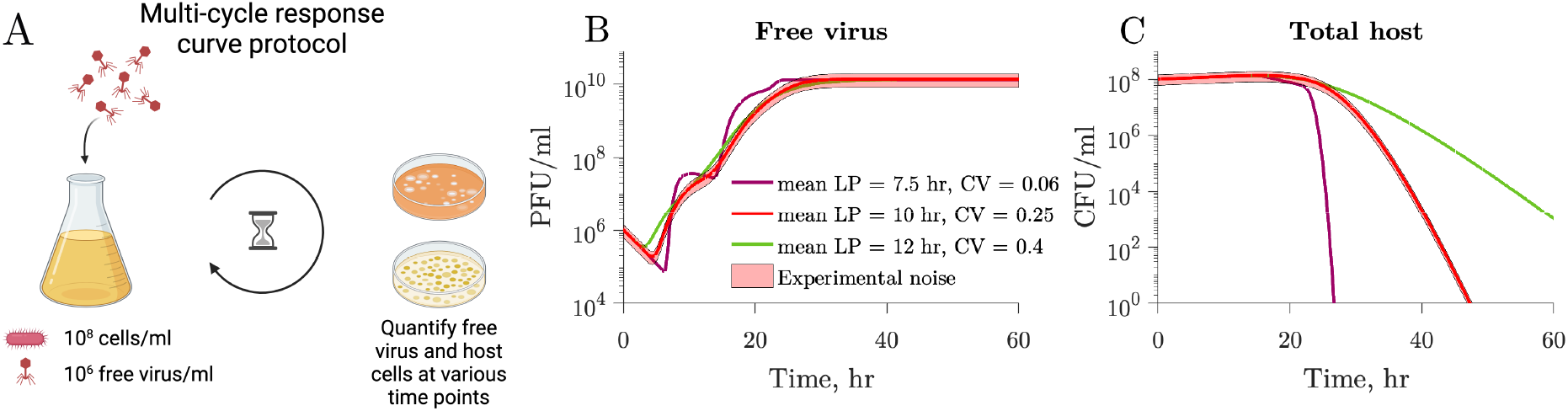
Multi-cycle response curves provide a better alternative for latent period distribution identification. **A)** We simulate an experimental protocol where infecting viral particles are added to a microbial population at MOI 0.01 (see Methods). Free-virus and host cells are quantified at multiple time points after infection. Unlike one-step growth curve protocols, there is no removal of free viral particles after an incubation period. The simulated time captures multiple rounds of infection. **B)** Free virus dynamics of three simulations with different latent period distributions but otherwise same viral and host parameters (Table 2). Note that multiple rounds of infection are observed. **C)** Corresponding host dynamics for the three simulations. While the one-step growth curves for the same parameters are highly similar (Figure 4), the multi-cycle response curves differ from each other.

### 2.3 Inferring latent period distributions from temporal dynamics

We developed a computational framework with the goal of inferring latent period distributions from multi-cycle response curves (Figure 6A). This framework involves fitting host and viral data to our population model of lytic infections (Equation 1), achieved by selecting combinations of model parameters that minimize the error between observed data and the model. The framework is comprised of two steps: (1) using a likelihood function to narrow the initial search space of parameters; (2) implementing a Bayesian Markov Chain Monte Carlo (MCMC) search. The Bayesian MCMC search is guided by prior distributions informed by the likelihood function used in the initial step (see Methods for further details). Using this approach, we can estimate host traits, *i.e*., growth rate (*μ*) and carrying capacity (*K*), and viral traits, *i.e*., adsorption rate (*ϕ*), burst size (*β*), and the latent period distribution (characterized by the mean *T* and CV) from host and viral dynamics data.

**Figure 6:**
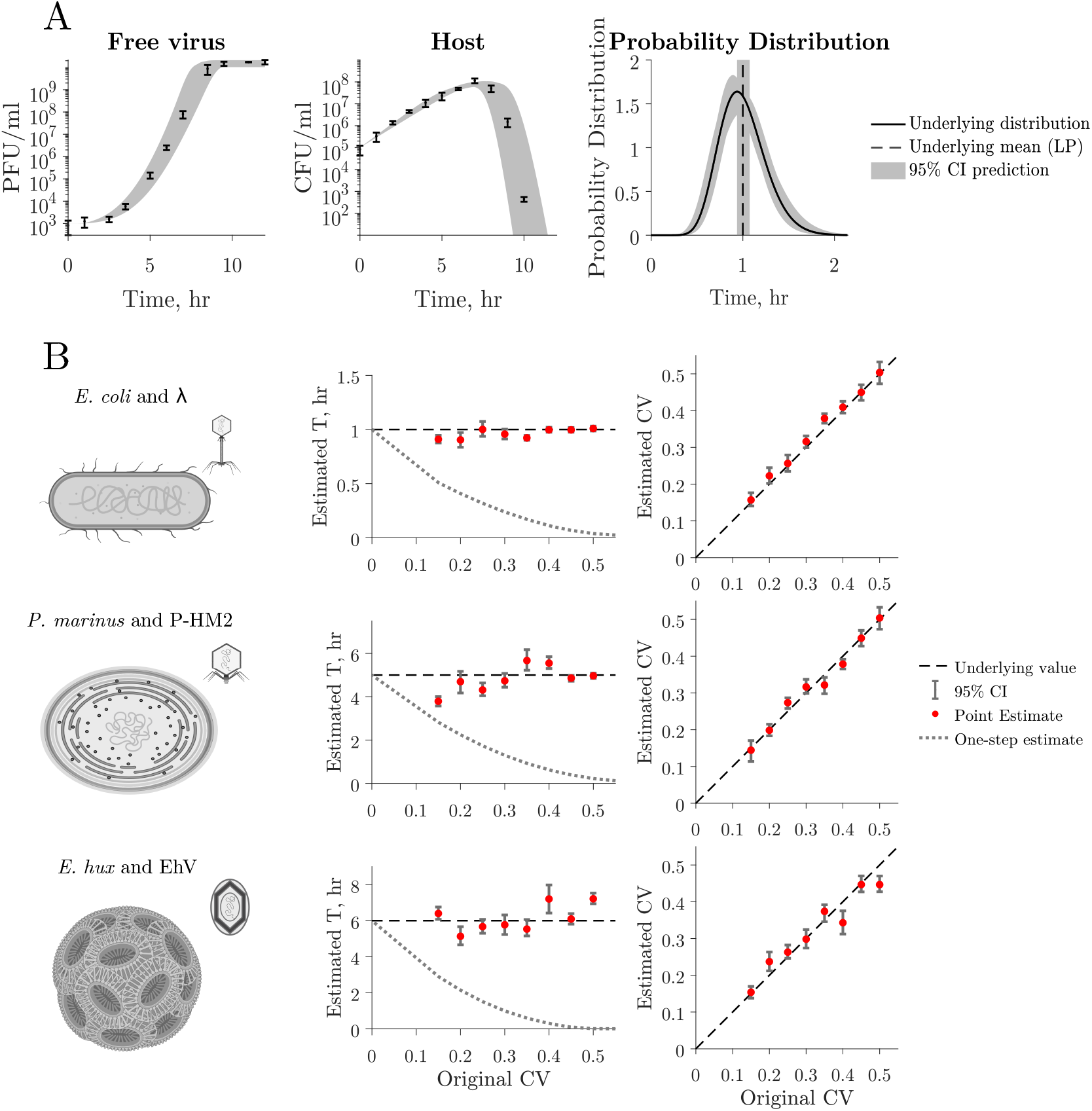
Latent period distribution estimated from simulated multi-cycle response curves. **A)** Estimation of the latent period distribution of a virus-microbe system by fitting a nonlinear dynamical model to the simulated time series with added noise. In this example, we estimate the latent period distribution of free-virus and host time series simulated data with added noise (left and middle panels). Thus, we have *a priori* knowledge of the underlying latent period distribution (black curve) of the system to evaluate our framework. We can accurately estimate the original mean (mean LP = 1 hr, black line) and CV = 0.25 and therefore estimate the original latent period distribution (compare black curve and confidence interval estimations). Gray shaded region indicates 95% confidence intervals. **B)** We use parameter values that capture the interactions of three biologically-relevant systems: *E. coli* and *λ* phage, Prochlorococcus and P-HM2, and *E. huxley* and EhV (Table 3). We model these systems assuming different latent period distribution dispersions. The dashed lines represent the original mean and CV values of the distribution used to create the data, red dots represent point estimates of the latent period mean (*T*) and CV, and error bars show 95% confidence intervals that fall within one order of magnitude of the original value across all simulations. The time of first burst obtained from the corresponding simulated one-step growth curves (dotted line) systematically underestimates the population mean, while our approach predicts the parameter value more accurately.

As our main interest is in inferring the latent period distribution, we test the accuracy of our approach at predicting these distributions using simulated data. Using simulated data allows us to have *a priori* knowledge of the underlying latent period distribution. We use our nonlinear dynamical model to generate host and viral time series. We sample the time series to obtain 10 data points equally spaced in time, consistent with current experimental standards, and add 30% normally distributed noise to mirror uncertainty in experimental measurements (Methods). To test our framework on a variety of biologically-relevant data, we generate dynamics using three different parameter sets that represent: 1) *E. coli* and *λ* phage, 2) the marine cyanobacteria *Prochlorococcus marinus* and the P-HM2 cyanophage, and 3) the eukaryotic microbe *Emiliania huxleyi* and a EhV (Table 3, Figure S3). We use literature point estimates of the latent period for the mean latent period (though Figure 2 suggests these are underestimates). The true, underlying CV of the latent period distributions for microbe-virus pairs is not known. Therefore, we generate data with CV-values ranging from 0.1 to 0.5, assessing our framework’s prediction accuracy across a large range of coefficients.

**Table 3:** Model parameter values for three distinct microbe-virus pairs. We simulate host and free virus temporal data using biologically relevant parameter values of three different microbe-virus pairs. Parameter values were recovered from [7] for the *E. coli* and *λ* and from [40] for the *Prochlorococcus* and P-HM2 cyanophage. References for *Emiliania huxleyi* and EhV parameters are shown next to the corresponding value.

| | *E.coli* and $\lambda$ | *P. marinus*<br>and P-HM2<br>cyanophage | *E. hux* and<br>EhV | |
| --- | --- | --- | --- | --- |
| Parameter | Value |  |  | Unit |
| $\mu$, growth rate | 1.2 | 0.035 | 0.015 [49] | hr<sup>-1</sup> |
| $K$, carrying capacity | $1 \times 10^8$ | $3 \times 10^9$ | $1 \times 10^9$ | CFU/ml |
| $\phi$, adsorption rate | $1 \times 10^{-8}$ | $9.3 \times 10^{-10}$ | $1.5 \times 10^{-7}$ [50] | ml/(CFU×hr) |
| $\beta$, burst size | 200 | 40 | 800 [51] | |
| $\eta$, lysis rate | 1 | 0.2 | 0.1667 [51] | hr<sup>-1</sup> |
| Initial condition | Value |  |  | Unit |
| $S_0$, initial microbial density | $10^5$ | $10^8$ | $10^5$ | CFU/ml |
| $V_0$, initial free virus density | $10^3$ | $10^6$ | $10^3$ | PFU/ml |

Consider first the viral population dynamics, host population dynamics, and latent period distribution associated with the *E.coli* and *λ* phage parameter set (Figure 6A). In this simulation, both free-virus and host density increase before host density starts to decay around 7 hours after initial inoculation. With a reduction in susceptible hosts available for infection, the free virus density starts to plateau shortly after. Note that these population dynamics reflect multiple mixed rounds of infection in contrast to the single cycle represented in one-step growth curves. The data fit is shown as the shadow region with a 95% confidence. The free virus and host data were simulated with an underlying distribution with latent period mean equal to 1 hr and CV of 0.25. The underlying distribution is correctly predicted at a 95% confidence. We observe that for this example, the individual parameters that describe the underlying latent period distribution (*T* and CV) fall within the 95% confidence interval for the predicted parameters (denoted by arrows in Figure 6B). Furthermore, the estimation for the average latent period mean is closer to the underlying value than that predicted by the corresponding one-step growth curve simulation (dotted line in Figure 6B).

The latent period mean and CV values predicted using our framework are close to those with which the data was generated for all simulations and all parameter sets for *E. coli* and *λ* phage, cyanobacteria and cyanophage, and Ehux and its associated giant virus (Figure 6B). Therefore, the underlying latent period distribution with which data was generated is recapitulated by the data fitting framework. Our estimates for latent period mean are closer to the population’s true underlying latent period than the time of first burst obtained from simulated one-step growth curves (dotted lines). In addition, we successfully predict all other host and viral traits (Figure S4). Hence, by incorporating cellular-level variability and fitting models to host and viral abundance data, our approach yields accurate estimates of the latent period distribution and other host and viral traits, for a variety of biologically-relevant systems.

## 3 Discussion

Here, we explored how individual-level variation in the latent period impacts virus-host dynamics and its consequences on the inference of viral life history traits, including the mean and variability of the latent period distribution. To do so, we developed a model of virus-host interactions that incorporates latent period variability. We show that latent period estimates obtained from current viral life-history trait estimation protocols – specifically the one-step growth curve – can deviate systematically and substantially from the actual mean latent period. The rationale is that variability in the latent period can lead to a subpopulation of infected cells lysing early, which is then misinterpreted as a shorter mean latent period for the population as a whole. Instead, we propose using host and viral dynamics to infer the population latent period via a ‘multi-cycle response curve’ protocol. By fitting nonlinear population models, we show that it is possible to recover unbiased estimates of the mean and variability of latent periods along with accurate estimates of other viral and cellular life history traits.

The present approach extends efforts leveraging model fitting to characterize viral life history traits [12, 39–42]. Similar to the modeling structure proposed here, other models of virus-host dynamics have incorporated explicit treatment of multiple infection compartments as an approximation to the delay between infection and lysis [12, 39, 43]. In these cases, the model interpretation does not link variability in the latent period with impacts on the estimate of the latent period itself. Inaccurate estimates of viral latent periods can lead to incorrect assumptions about virus turnover, potentially leading to over-estimates of viral-induced mortality at population scales. Instead, accurate and unbiased estimates of viral traits are required to inform ecological and ecosystem models [7, 42, 44, 45].

The use of multi-cycle response curves leverages information in both viral and host population time series to connect cellular-level processes to population-level dynamics. In doing so, early increases in virus population dynamics are linked to the early components of variable latent period distributions rather than an indication of a systematically faster lytic process. This feature is likely relevant across virus-host systems. However, more work is needed to identify the mechanisms underlying variability, including those that impact other viral traits. For example, while correlations between latent period and burst size have been observed, with phage strains with longer latent periods having larger burst sizes [13], recent studies suggest that variation in (induced) lysis time does not contribute to observed burst size variability [46, 47]. Caution in the interpretation of multi-cycle response curves is required, as viral traits often depend on host growth rate which can vary in experimental timescales [12]. In addition, the current method could be extended in multiple ways, *e.g*., to include virion decay on the timescale of the protocol, which may be relevant for extending the method *in situ* and/or *in vivo* where reduced viral stability is expected or to include coinfection of a single cell by multiple viruses.

In summary, for more than 80 years, the one-step growth curve has been the standard to characterize essential viral life-history traits: the latent period and burst size [15]. As we have shown, this protocol comes with a significant caveat: leading to estimates of more rapid lysis when neglecting the impacts of latent period variability. Our work suggests that reframing the latent period as a distribution is essential to improve viral life-history trait estimates and to connect viral traits to population-level dynamics. In practice, the use of multi-cycle response curves leverages host and viral time series to reduce biases in trait inferences. Moving forward, we are optimistic that embedding measurements of both viral and host time series in the context of mathematical models with explicit treatment of variability can be used to improve inference in experiments and estimates of viral traits in environmental and therapeutic contexts.

## 4. Methods

### 4.1 Latent period distribution incorporated into a nonlinear population model of lytic viral infections

We use a coupled system of nonlinear differential equations to model virus-host dynamics, including susceptible cells, *S*, free viruses, *V*, exposed cells, *E*, and actively-infected cells, *I*. In this model, susceptible host cells, *S*, are infected by free virus particles, *V*. We assume that viruses and hosts are well-mixed. Under these assumptions, the incidence, the rate at which susceptible cells are infected, is given by the mass action term: *i*(*t*) = *ϕ S V*, where *ϕ*(ml/hr) denotes the adsorption rate.

To incorporate variability in latent period, we assume that before entering the actively-infected stage, *I*, infected cells advance through several exposed *E* stages: *E*_1_, …, *E*_*n*_, where *n* is a non-negative integer; infected cells move between stages at a rate of (*n* + 1) *η* with exponentially distributed times, where *η* is the lysis rate. As the cells remain in each stage for a period of *T/*(*n* + 1) on average and there are *n* + 1 stage transitions, the average time from adsorption (*i.e*., entering the first exposed class, *E*_1_) to cell burst (*i.e*., exiting the actively-infected class, *I*) is the latent period mean, equal to the inverse of the mean lysis rate, *T* = 1*/η*. At the end of the actively-infected stage, *I*, the cell bursts and free virus, *V*, increases as a result of viral release of *β* virions. The system of nonlinear, ordinary differential equations can be written in the form:

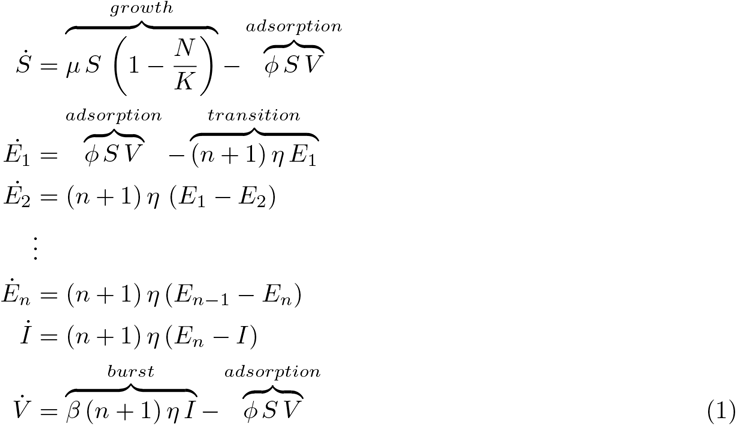

where *μ* (1/hr) denotes the maximal cellular growth rate, *K* (1/ml) denotes the cellular carrying capacity in the absence of viruses, and 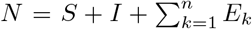 is the total cell population. Note that when the number of *E* stages *n* equals 0, the model is reduced to the SIV model where *İ* = *i*(*t*) *− η I*. We assume that infected cells at any stage of infection do not grow and that cell death rates and virus washout rates are negligible compared to other key rate constants of the system. See Table 1 for model parameter descriptions. Hence, this model describes the latent period distribution as an Erlang distribution with shape *n* + 1, the number of exposed (*E*) compartments plus the infected (*I*) compartment, and rate *η*, the lysis rate. In this form, the mean (*T*), variance (*σ*^2^), and coefficient of variation (*σ/T*) of the latent period are given by:

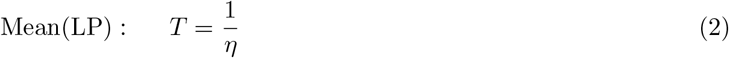

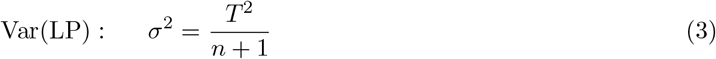

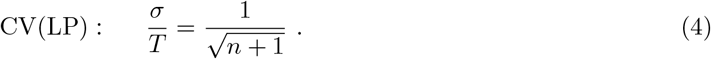

The number of *E* compartments modulates the dispersion of the distribution through the coefficient of variation (CV), with larger *n* resulting in tighter distributions with smaller CV (Figure S1). Note that requiring *n* to be an integer limits the CV values that can be simulated. For example, *n* = 0 corresponds to *CV* = 1, while *n* = 1 corresponds to *CV ≈* 0.7, meaning that CV values between 0.7 and 1 cannot be represented using our model. Latent period distributions with CV lower than 0.5 can be simulated with our model at a tolerance of 0.05. Based on lysis timing variability of induced lysogens [35], we expect latent period CV values in natural systems to be *<*0.5, consistent with our model’s effective coverage of latent period variability.

### 4.2 Model implementation

#### 4.2.1 One-step growth curve

When using our model to simulate one-step growth curves, we replicate the protocol in experimental one-step growth curves by simulating the addition of phage to a microbial population in exponential growth phase. We initialize the system with *S* = 10^8^ CFU/ml and *V* = 10^6^ PFU/ml with MOI 0.01. We dilute the system 1000-fold 10 minutes after phage addition to reduce microbe-virus encounters and prevent further adsorption [15, 16]. After dilution, free virus (*V*) is quantified at multiple timepoints (Figure 2A). Parameters used to simulate the curves in Figure 2 can be found in Table 1. Parameters used to simulate one-step growth curves in Figure 4 can be found in Table 2. Errors in Figure 5 were calculated using 200 equally-spaced time points.

#### 4.2.2 Multi-cycle response curve

In our simulation of multi-cycle response curves, we initialize the system with *S* = 10^8^ CFU/ml and *V* = 10^6^ PFU/ml with MOI 0.01. Free virus (*V*) and total host 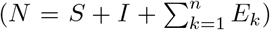 are sampled at multiple time points. In contrast to the one-step growth curve simulations, there are no incubation period and subsequent dilution. The experiment time is longer in multi-cycle response curve than one-step growth curve simulations to capture multiple rounds of infection. The parameter values used to simulate temporal dynamics in Figure 5 are the same as for the one-step growth curve simulations in Figure 4 and can be found in Table 2.

To simulate the dynamics of a variety of host and viral systems, we use reference parameter values for three different microbe-virus pairs: *E. coli* and *λ* phage, *Emiliania huxleyi* and EhV, and *Procholorococcus* and P-HM2 cyanophage (Table 3). Different latent period distributions are generated by fixing the mean (Equation 2) and varying the CV (Equation 4) from 0.15 to 0.5 (Figure S3). For every set of parameters, we model two scenarios: a control where the host grows in the absence of virus, and an experiment with free-virus and an initially susceptible microbial population. The control simulation is used to estimate host-only parameters. The corresponding initial microbial and free-virus densities are presented in Table 3. We simulate three experimental replicates by sampling the dynamics at 10 equally spaced time points and adding measurement noise according to a normal distribution with mean equal to 0 and standard deviation of 30% constrained to non-negative numbers. The standard deviation value was chosen from observations of experimental replicates [48].

### 4.3 Latent period distribution inference

We use a computational framework to estimate latent period distributions from host and free virus temporal dynamics data by fitting to our population model of lytic infections (Equation 1). Our framework predicts all host and viral associated parameters comprised in the model, *i.e*., host growth rate (*μ*), carrying capacity (*K*), adsorption rate (*ϕ*), burst size (*β*), and latent period distribution associated parameters (*T* and CV). The framework is comprised of two main steps: (1) a likelihood function to narrow the search space of parameters; (2) a Markov Chain Monte Carlo (MCMC) search with prior distributions informed by the likelihood function in step (1).

In step (1), rough parameter ranges are found using a grid search for the maximum likelihood parameter combination from a range of biologically plausible parameter values [52]. In step (2), we implement MCMC using the Turing package in Julia [53] and inform prior distributions using the predictions obtained in step (1). The resulting posteriors are then used as priors for a second round of MCMC. Details on the prior distributions and convergence analysis can be found in the Supplementary information. We obtain 95% confidence intervals by sampling the MCMC posterior distributions. We test our framework by fitting data generated with added noise, as explained above, for which the underlying latent period distribution is known.

### 4.4 Code availability

We implement the model (Equation 1) in Julia v1.7.2 [54] using the DifferentialEquations package v7.1 [55] and Matlab R2023b [56]. All code for simulations and plotting is available at: github.com/WeitzGroup/LatentPeriodVariability, and archived at: doi.org/10.5281/zenodo.11085440

## 5 Acknowledgments

The research effort was enabled by support from grants from the Simons Foundation (JSW, Award ID 722153), the Blaise Pascal Chair (JSW), the Chateaubriand Fellowship (MDM), and the National Council of Humanities, Science and Technology of Mexico (MDM). Funding sources had no role or influence on study design, analysis, interpretation, or submission. We thank Stephen Beckett and Adriana Lucia-Sanz for code review. This research was supported in part through research cyberinfrastructure resources and services provided by the Partnership for an Advanced Computing Environment (PACE) at the Georgia Institute of Technology, Atlanta, Georgia, USA. Illustrations were created with BioRender.com.

