## Supplementary Material for "Accounting for Cellular-Level Variation in Lysis: Implications for Virus-Host Dynamics"

### S1 Microbe-virus model and Parameter inference

To facilitate the reading of the supplementary material, we include the model description found in the main text.

#### S1.1 Nonlinear population model of lytic phage infections

We use a coupled system of nonlinear differential equations to model virus-host dynamics, including susceptible cells, $S$, free viruses, $V$, exposed cells, $E$, and actively-infected cells, $I$. In this model, susceptible host cells, $S$, are infected by free virus particles, $V$. We assume that viruses and hosts are well-mixed. Under these assumptions, the incidence, the rate at which susceptible cells are infected, is given by the mass action term: $i(t) = \phi S V$, where $\phi$(ml/hr) denotes the adsorption rate.

To incorporate variability in latent period, we assume that before entering the actively-infected stage, $I$, infected cells advance through several exposed $E$ stages: $E_1, \ldots, E_n$, where $n$ is a non-negative integer; infected cells move between stages at a rate of $(n+1)\eta$ with exponentially distributed times, where $\eta$ is the lysis rate. As the cells remain in each stage for a period of $T/(n+1)$ on average and there are $n+1$ stage transitions, the average time from adsorption (*i.e.*, entering the first exposed class, $E_1$) to cell burst (*i.e.*, exiting the actively-infected class, $I$) is the latent period mean, equal to the inverse of the mean lysis rate, $T = 1/\eta$. At the end of the actively-infected stage, $I$, the cell bursts and free virus, $V$, increases as a result of viral release of $\beta$ virions. The system of nonlinear, ordinary differential equations can be written in the form:

$$
\begin{aligned}
\dot{S} &= \overbrace{\mu\, S\left(1-\frac{N}{K}\right)}^{growth} - \overbrace{\phi\, S\, V}^{adsorption} \\
\dot{E}_1 &= \overbrace{\phi\, S\, V}^{adsorption} - \overbrace{(n+1)\,\eta\, E_1}^{transition} \\
\dot{E}_2 &= (n+1)\,\eta\,(E_1 - E_2) \\
&\vdots \\
\dot{E}_n &= (n+1)\,\eta\,(E_{n-1} - E_n) \\
\dot{I} &= (n+1)\,\eta\,(E_n - I) \\
\dot{V} &= \overbrace{\beta\,(n+1)\,\eta\, I}^{burst} - \overbrace{\phi\, S\, V}^{adsorption}
\end{aligned} \tag{1}
$$

where $\mu$ (1/hr) denotes the maximal cellular growth rate, $K$ (1/ml) denotes the cellular carrying capacity in the absence of viruses, and $N = S + I + \sum_{k=1}^{n} E_k$ is the total cell population. Note that when the number of $E$ stages $n$ equals 0, the model is reduced to the SIV model where $\dot{I} = i(t) - \eta\, I$. We assume that infected cells at any stage of infection do not grow and that cell death rates and virus washout rates are negligible compared to other key rate constants of the system. See Table 1 for model parameter descriptions. Hence, this model describes the latent period distribution as an Erlang distribution with shape $n + 1$, the number of exposed ($E$) compartments plus the infected ($I$) compartment, and rate $\eta$, the lysis rate. In this form, the mean ($T$), variance ($\sigma^2$), and coefficient of variation ($\sigma/T$) of the latent period are given by:

$$
\text{Mean(LP)}: \quad T = \frac{1}{\eta} \tag{2}
$$

$$
\text{Var(LP)}: \quad \sigma^2 = \frac{T^2}{n+1} \tag{3}
$$

$$
\text{CV(LP)}: \quad \frac{\sigma}{T} = \frac{1}{\sqrt{n+1}}\,. \tag{4}
$$

The number of $E$ compartments modulates the dispersion of the distribution through the coefficient of variation (CV), with larger $n$ resulting in tighter distributions with smaller CV (Figure S1). Note that requiring $n$ to be an integer limits the CV values that can be simulated. For example, $n = 0$ corresponds to $CV = 1$, while $n = 1$ corresponds to $CV \approx 0.7$, meaning that CV values between 0.7 and 1 cannot be represented using our model. Latent period distributions with CV lower than 0.5 can be simulated with our model at a tolerance of 0.05. Based on lysis timing variability of induced lysogens [1], we expect latent period CV values in natural systems to be <0.5, consistent with our model's effective coverage of latent period variability.

### S1.2 Parameter inference

This section provides extended details for the parameter estimation section in the Methods of the main text. All code is written in Julia v1.8 [2]. Our computational framework fits host and viral temporal data to our nonlinear population model of lytic infections to recover the parameters that allow the model to better represent the data.

First, host-only parameters, *i.e.*, host growth rate ($\mu$) and carrying capacity ($K$) are estimated using temporal data of host without added phage that captures the exponential and stationary phases. The framework is composed of two steps: (1) The log-transformed data is segmented into two lines, one representing the slope of the exponential phase and the other one representing the growth plateau. The slope of the exponential phase is assumed to approximate the host growth rate ($\mu$) and the $y$-intercept of the stationary phase is assumed to approximate the carrying capacity ($K$); (2) The point estimates obtained in step (1) are used to inform priors for a Markov Chain Monte Carlo (MCMC) search. Details on the MCMC search implementation are provided below. These host-only predicted parameter values are used in the following steps of the parameter inference.

Second, the computational framework estimates the model microbe-virus parameters, *i.e.*, adsorption rate ($\phi$), burst size ($\beta$), lysis rate ($\eta$), and the number of E compartments ($n$) using host and free virus temporal data from 'multi-cycle response curves' where free virus is added to a microbial population and both free virus ($V$) and total host ($N$) densities are measured at multiple time points. The framework's main aim is to recapitulate the latent period distribution described by the latent period mean ($T = 1/\eta$) and coefficient of variation ($\text{CV} = (n+1)^{1/2}$). The framework for microbe-virus trait estimation is comprised of two main steps: (1) a likelihood function to narrow the search space for all microbe-virus parameters; (2) a Markov Chain Monte Carlo (MCMC) search with prior distributions informed by the likelihood function in step (1). In step (1), rough parameter ranges are found using a grid search for the maximum likelihood estimate (MLE) of the parameter combination. In step (2), we implement MCMC using confidence intervals obtained in step (1) to inform prior distributions. As chains can converge to different parameter ranges, we choose the set of chains that converge to the same values for all parameters and that minimize the error chains. The resulting posteriors are then used as priors for a second round of MCMC. We obtain 95% confidence intervals by sampling the resulting posterior distributions.

#### S1.2.1 MCMC implementation

For step (2) in parameter estimation of both host-only and microbe-viral parameters we use Markov Chain Monte Carlo (MCMC) implemented in the probabilistic inference package Turing [3] in the Julia language v1.8 [2]. MCMC is a class of algorithms that aim to obtain the equilibrium probability distribution of the model parameters [4]. We use the No-U-Turn Sampler (NUTS) from the Turing package [3] to sample the posterior distribution. For each implementation, we ran MCMC chains for 2000 iterations with a 1000-iteration warm-up period. The target acceptance ratio was set to 0.45. We obtained high autocorrelation for the cases where the coefficient of variation is small. In these cases, we achieved chain convergence by thinning chains to obtain 3000 steps.

#### S1.2.2 Chain convergence analysis

To ensure the convergence of the MCMC chains in step (2), we calculated the potential scale reduction, $\hat{R}$ [5] for the pooled 10000-iteration long chains. For each parameter, $\hat{R}$ measures the ratio of the average variance in samples of the chain to the variance of the whole chain. When the chains have converged around a posterior distribution, $\hat{R}$ will be centered around 1. If the chains have not converged to a single distribution, $\hat{R}$ values will be decimals larger than 1. The $\hat{R}$ calculated for all our chains is centered below 1.1, suggesting convergence.

We computed the Effective Sample Size (ESS, $N_{eff}$) ratio as $N_{eff}$ divided by the number of iterations in the pooled chain ($N = 3000$). A low ratio, an empirical common cutoff of $\frac{N_{eff}}{N} << 0.1$, suggests sampling issues and unreliability to estimate confidence intervals. The ESS ratio for all our chains is close to 0.1. Both $\hat{R}$ and $N_{eff}$ were calculated with the `ess_rhat` function from the `MCMCDiagnosticTools` Julia package [6] (Figure S5).

#### S1.2.3 Parameter ranges and MCMC priors

For step (1) of the microbe-virus parameter inference, we do a Maximum Likelihood grid search with biologically relevant parameter value ranges. The prior distributions for the MCMC search in step (2) are informed by estimations obtained from step (1). The prior distributions are given by a $LogNormal(\theta, \nu)$, where $\theta = \ln mode + \frac{2}{3}\ln\frac{mean}{mode}$ and $\nu = \sqrt{\frac{2}{3}\ln\frac{mean}{mode}}$. For the first round of MCMC the *mode* of the distribution is equal to the Maximum Likelihood point estimates obtained in step (1), and the distribution *mean* is equal to two times the point estimate. For the second round of MCMC (only used for microbe-virus parameters), the *mode* is equal to the mean of the parameter chain from the previous round, and the distribution *mean* equals the *mode* plus two times the chain's standard deviation.

| Host species | Virus name | Reported Latent Period | Source | Figure |
|---|---|---|---|---|
| *S. eneterica* albany | vB_SalP_TR2 | 15 min | Shang, 2021 [7] | 2B |
| *A. baumannii* AB1 | Abp1 | 15 min | Huang, 2013 [8] | 3 |
| *P. syringae* pv. actinidiae strain XWY0007 | $\phi$XWY0013 | 20 min | Yin, 2019 [9] | 2 (squares) |
| *S. aureus* MRSA t127/4 | UPMK_1 | 20 min | Dakheel, 2019 [10] | 5A |
| *E.coli* strain CR63 | RB69_WT | 21 min | Abedon, 2003 [11] | 2B (solid squares) |
| *A. hydrophila* strain 4572 | AhyVDH1 | 50 min | Cheng, 2021 [12] | 2B |
| *Vibrio spp.* strain B8D | P8D | 60 min | Yu, 2013 [13] | 3D |
| *S. maltophilia* strain T39 | $\phi$SMA5 | 60 min | Chang, 2005 [14] | 2 |
| *S. marcescens* strain wk2050 | vB_SmaA_2050H1 | 80 min | Tian, 2019 [15] | 3A |
| *P. syringae* strain CRA-FRU8.43 | $\phi$PSA1 | 100 min | Di Lallo, 2014 [16] | 3 (squares) |
| *D. shibae* strain DFL12 | DS-1410Ws-06 | 2 hr | Li, 2016 [17] | 2C (open circles) |
| *Synechococcus spp.* strain WH7803 | Syn9 | 4 hr | Doron, 2015 [18] | 1B (blue) |

Table S1: **Data sources for Figure 1** Sources for one-step growth curves shown in Figure 1 including host species with strains and virus used to perform the experiments. The latent period is explicitly reported in the text and/or figure captions of the sources.

| Strain name in [19] | Strain name in [1] | Population-level measurement (min) Lysis time ± sd (r) [19] | Cellular-level measurement (min) Mean lysis time ± sd (n) [1] | CV [1] |
|---|---|---|---|---|
| $S$ | IN56 (WT) | 51.0 ± 0.92 (3) | 65.1 ± 3.24 (230) | 0.05 |
| $S105$ | IN62 | 42.7 ± 0.54 (3) | 54.3 ± 3.42 (136) | 0.063 |
| $S105_{C51S}$ | IN61 | 35.0 ± 0.92 (3) | 45.7 ± 2.92 (274) | 0.064 |
| $S_{S68C}$ | IN66 | 63.0 ± 1.60 (3) | 82.2 ± 5.87 (189) | 0.071 |
| $S_{C51S/F94C}$ | IN64 | 42.7 ± 0.54 (3) | 48.4 ± 4.60 (63) | 0.095 |
| $S105_{C51S/L14C}$ | IN68 | 38.5 ± 1.80 (4) | 54.1 ± 5.14 (153) | 0.095 |
| $S_{C51S/L14C}$ | IN69 | 38.5 ± 1.80 (4) | 45.0 ± 4.38 (119) | 0.097 |
| $S105_{C51S/S76C}$ | IN63 | 28.3 ± 1.06 (3) | 41.2 ± 4.55 (209) | 0.11 |
| $S105_{C51S/I13C}$ | IN67 | 37.5 ± 0.94 (4) | 57.6 ± 6.71 (212) | 0.116 |

Table S2: **Lysis time in $\lambda$ phage lysogens is consistently underestimated when using population-level protocols.** The lysis time upon thermal induction of isogenic $\lambda$ strains that differ only in mutations in the $S$ gene, which encodes a holin-antiholin system that modulates lysis timing, was determined using turbidity assays in [19]. In a subsequent study, the lysis time of individual cells was measured for the same strains using microscope tracking [1]. The same induction protocol was followed in both studies. Notably, the mean lysis time calculated from cellular-level measurements is consistently longer than the one obtained from population-level protocols. *S105* strains have a mutation that abolishes antiholin expression. (r) represents the number of replicates in turbidity assays. (n) represents the number of individual cells observed and may represent the pool of multiple experiments.

| Parameter | Parameter description | Step (1) Search space | Step (2) Prior distribution* | Step(2) Prior bounds |
|---|---|---|---|---|
| $\mu$ | Growth rate | NA | $LogNormal(\theta,\nu)$ | [0, 2] |
| K | Carrying capacity | NA | $LogNormal(\theta,\nu)$ | $[10^3, 10^{10}]$ |
| $\phi$ | Adsorption rate | $[10^{-12}, 10^{-6}]$ | $LogNormal(\theta,\nu)$ | $[10^{-11}, 10^{-5}]$ |
| $\beta$ | Burst size | [1,1000] | $LogNormal(\theta,\nu)$ | [1, 1000] |
| $\eta$ | Lysis rate | [0.025, 12] | $LogNormal(\theta,\nu)$ | [0.083, 6] |
| CV | Coefficient of variation | [0.05, 0.5] | $LogNormal(\theta,\nu)$ | [0.03, 1] |
| $\sigma_{loglikelihood,v}$ | Error chain for viral series | NA | $InvGamma(1, 0.5)$ | [0, 2] |
| $\sigma_{loglikelihood,h}$ | Error chain for host series | NA | $InvGamma(1, 0.5)$ | [0, 2] |

Table S3: **Parameter ranges and MCMC priors used for parameter inference.** The specific values for $\theta$ and $\nu$ are described in the text above.

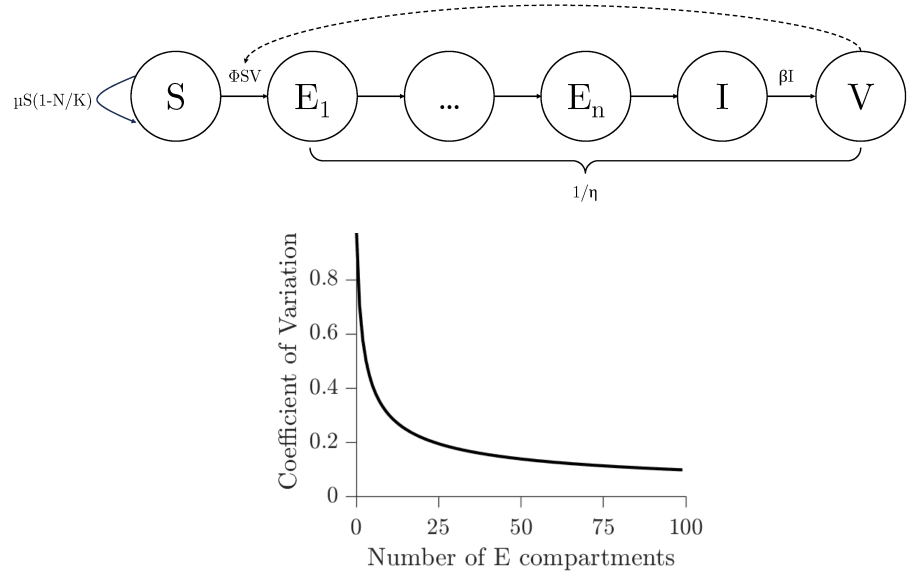


Figure S1: **Latent period variability incorporated in a lytic infection model. A)** We construct a compartmental model of lytic viral infections. Susceptible cells, $S$, grow in a logistic manner determined by the growth rate ($\mu$) and carrying capacity ($K$). Susceptible cells are infected by free viral particles, $V$, dependent on the susceptible cell, viral density, and the adsorption rate ($\phi$). Before entering the actively infected stage, $I$, infected cells advance through several exposed $E$ stages. The number of $E$ stages ($n$) is a parameter of the model that determines the coefficient of variation (CV) of the latent period distribution. After reaching the actively infected stage $I$, cells burst yielding viral offspring with burst size $\beta$. The average time before cell burst is the inverse of the lysis rate ($\eta$). **B)** The number of E compartments ($n$) is treated as a model parameter. The CV of the distribution scales like $(n+1)^{-1/2}$.

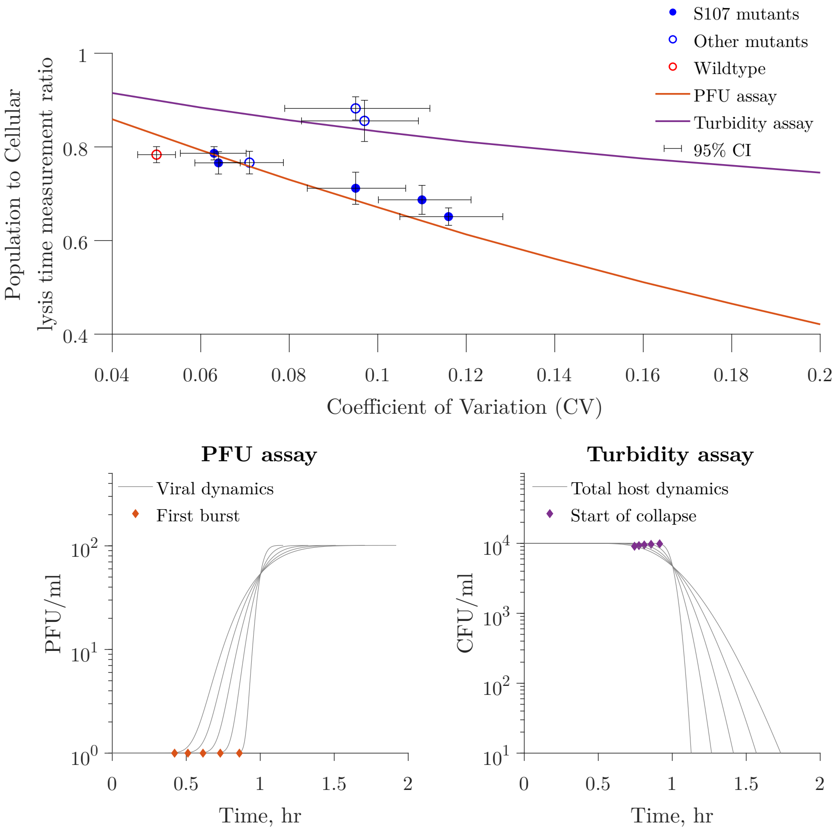


Figure S2: **Lysis time estimation bias in $\lambda$ phage lysogens using population-level protocols is dependent on the coefficient of variation.** Top, the population to cellular-level lysis time measurement ratio decreases with larger CV, as a result of early lysis events. Confidence intervals were calculated by multiple resampling using experimental mean and standard deviation assuming normality, as explained in [20]. Our model predicts a similar trend where the mean lysis time underestimation increases with CV when using either PFU assays (represented by the time of first visible burst, bottom left panel) or turbidity assays (represented by the start of host population collapse, bottom right panel) as indicators of lysis time. This trend is parameter-free, as we simulate a perfectly synchronized population of infected cells by initializing the simulations with the total host density in the first infection compartment ($E_1$). Given that infected cells are assumed to not grow, host-only parameters (growth rate, $\mu$ and carrying capacity, K) are irrelevant. The time of first burst and start of host population collapse can be characterized independent of burst size ($\beta$). Our model also suggest that assays using host dynamics may estimate the mean lysis time more accurately than PFU-based methods (compare purple to orange lines in top panel).

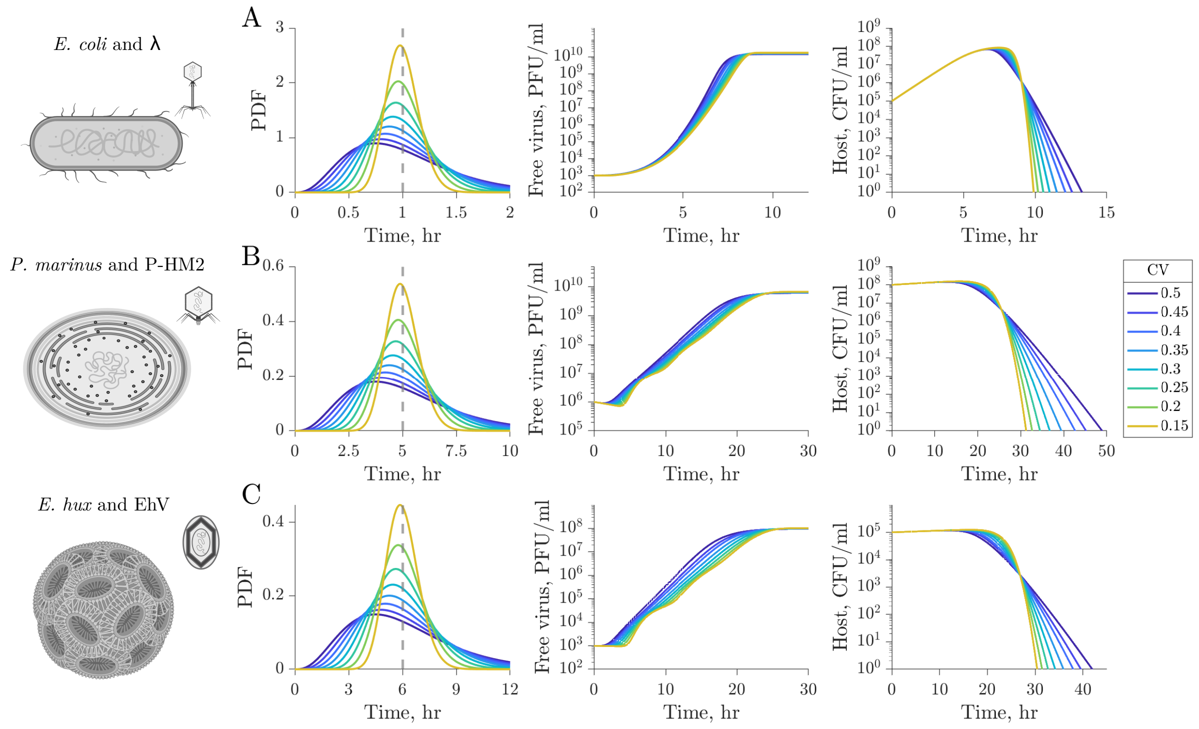


Figure S3: ***In silico* multi-cycle response curves of biological systems.** Using our virus-microbe dynamical model (see Methods) we simulate the dynamics of three different virus-microbe pairs. Parameter values were obtained from literature estimations of the host and virus life-history traits (Table 3). Multi-cycle response curves were simulated for the different systems varying the latent period coefficient of variation to range between 0.05 and 0.5 (distributions are shown on the left). The free virus and host densities are shown in the middle and right panels, respectively.

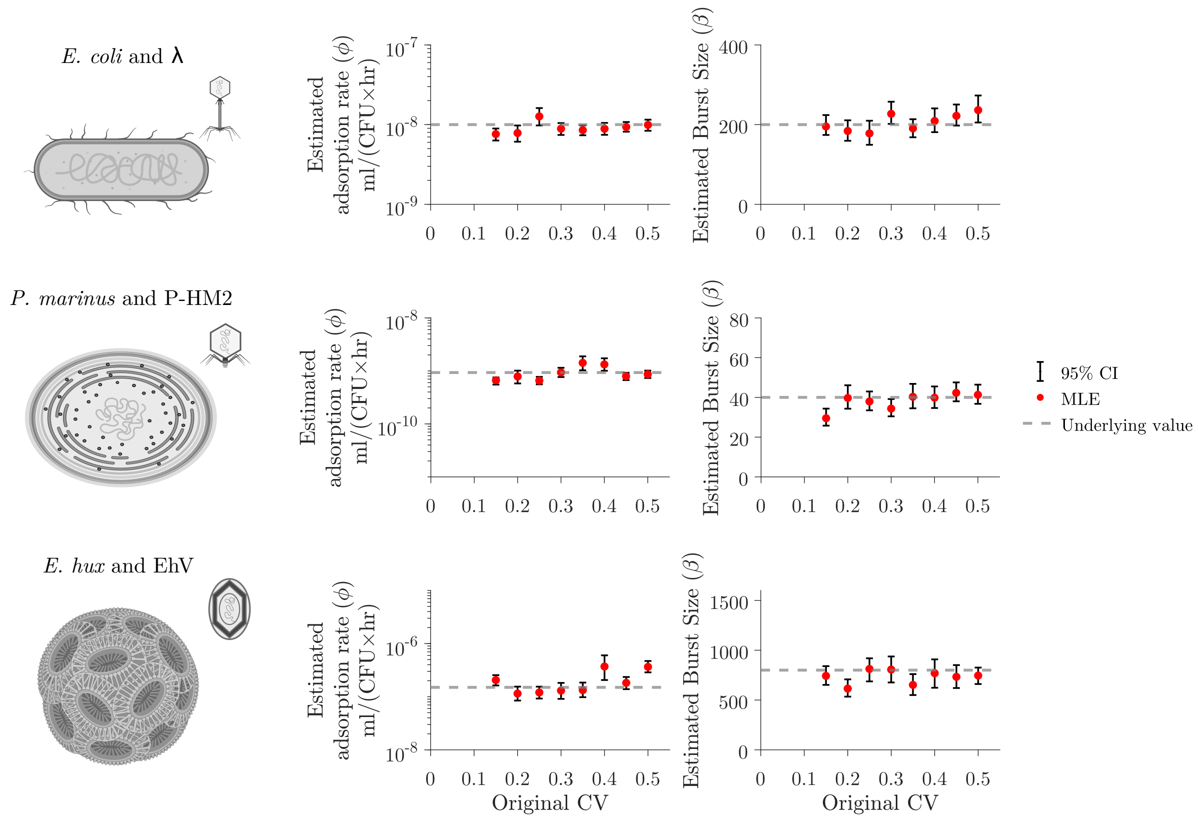


Figure S4: **Prediction of non-latent period virus-host traits.** We use parameter values that capture the interactions of three biologically-relevant systems: *E. coli* and $\lambda$ phage, Prochlorococcus and P-HM2, and *E. huxley* and EhV (Table 3). We model these systems assuming different latent period distribution dispersions. The dashed lines represent the original adsorption rate and burst size values used to create the data. We use a free virus and host data-model fitting approach to predict viral and host traits. Red dots represent point estimates of the parameters, and error bars show 95% confidence intervals that fall within one order of magnitude of the original value across all experiments.

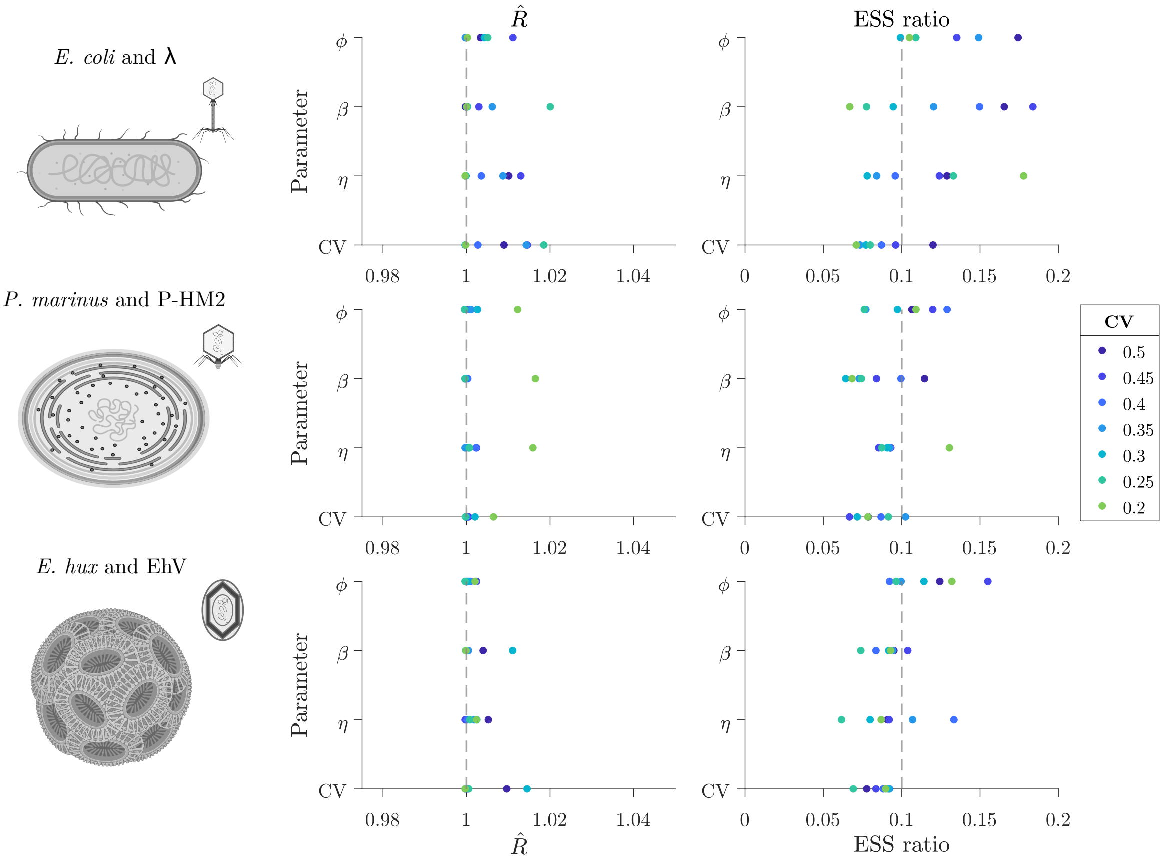


Figure S5: **Chain convergence analysis.** We calculated the potential scale reduction, $\hat{R}$ (left panels) and the Effective Sample Size ratio, ESS ratio(rigth panels) of our final MCMC chains. $\hat{R}$ measures the ratio of the average variance in samples of the chain to the variance of the whole chain. If the chains have not converged to a single distribution, $\hat{R}$ values will be decimals larger than 1. The Effective Sample Size rato is the the Effective Sample Size divided by the number of iterations in the pooled chain. A low ratio, an empirical common cutoff of $\frac{N_{eff}}{N} << 0.1$, suggests sampling issues and unreliability to estimate confidence intervals.
